# Secreted cytokines provide local immune tolerance for human stem cell-derived islets

**DOI:** 10.1101/2022.05.09.487072

**Authors:** Dario Gerace, Quan Zhou, Jennifer Hyoje-Ryu Kenty, Elad Sintov, Xi Wang, Kyle R Boulanger, Hongfei Li, Douglas A Melton

## Abstract

Immunological protection of transplanted stem cell-derived islet (SC-islet) cells is yet to be achieved without chronic immunosuppression or encapsulation. Existing genetic engineering approaches to produce hypoimmunogenic SC-islet cells have so far shown variable results. Here, we show that targeting the human leukocyte antigens (HLAs) and PD-L1 alone do not sufficiently protect SC-islet cells from xeno- or allo-rejection. As an addition to these approaches, we genetically engineered SC-islet cells to secrete the cytokines IL-10, TGF-β and modified IL-2 such that they promote a tolerogenic local microenvironment by activating and expanding regulatory T cells (T_regs_). These cytokine-secreting human SC-islet cells prevented xeno-rejection for up to 9 weeks post-transplantation in B6/albino mice. Thus, hESCs engineered to induce a tolerogenic local microenvironment may represent a source of replacement SC-islet cells that do not require encapsulation or immunosuppression for diabetes cell replacement therapy.

## Introduction

T1D is an autoimmune disease that results in the destruction of the insulin-producing beta cells of the pancreas (Atkinson and Maclaren, 1994). Cadaveric whole pancreas or islet transplantation are successful treatments for T1D; however, these are hampered by the limited number of donors and the requirement for lifelong immunosuppression (Shapiro et al., 2006). To address the shortage of islet material, several protocols have been developed to steer the *in vitro* differentiation of human induced pluripotent stem cells (iPSCs) into functional SC-islets that include glucose-response beta cells (D’Amour et al., 2006; Millman et al., 2016; Nair et al., 2019; Pagliuca et al., 2014; Rezania et al., 2014; Russ et al., 2015; Veres et al., 2019).

Current strategies to protect allografted islet cells include encapsulation (Alagpulinsa et al., 2019; Bochenek et al., 2018), modifying the patient’s immune system by co-administration of biologicals such as low dose IL-2 and anti-CD3 (Tepluzimab) (Hartemann et al., 2013; Herold et al., 2019), and/or genetically modifying the SC-islets. Since the main contributors of immune recognition and rejection are the human leukocyte antigens (HLAs), targeting the HLAs has been performed in iPSCs to reduce or eliminate the immune response against foreign cells (Castro-Gutierrez et al., 2021; Deuse et al., 2019; Gornalusse et al., 2017; Han et al., 2019; Harding et al., 2019; Riolobos et al., 2013; Xu et al., 2019; Yoshihara et al., 2020). To date, the engineering strategies demonstrating some immune-protection of SC-islet cells have employed lentiviral over-expression of PD-L1 and the selective retention of a single HLA-A2 allele in HLA-B/C deficient cells (Parent et al., 2021; Yoshihara et al., 2020). However, lentiviral transgene over-expression is limited by transgene silencing (Herbst et al., 2012; Wen et al., 2021), and the retention of a single HLA-A2 allele on SC-islet cells may result in recurrent autoimmune rejection mediated by antigen-specific and tissue-resident memory T cells (Abou-Daya et al., 2021; Monti et al., 2008; Stegall et al., 1996).

To address the issue of transgene silencing during differentiation into SC-islet cells, we employ a transgene targeting strategy that makes use of the constitutively expressed GAPDH gene in primary human islets and SC-islet cells (Gerace et al., 2021; Sintov et al., 2021). We generate hypoimmunogenic SC-islet cells that over-express PD-L1 and the HLA-E long-chain fusion in both HLA-competent and deficient settings. In these studies, PD-L1 over-expression did not protect SC-islet cells from xeno-rejection and over-expression of the HLA-E single-chain fusion was not required to inhibit primary human NK cells. In addition to immune evasion, we also address immune modulation and show that constitutive secretion IL-10, TGF-β and the IL-2 mutein does not impair SC-islet cell function *in vitro* and does provide protection of SC-islet cells from xeno-rejection for up to 9 weeks after transplantation. Thus, these immune-modulating SC-islet cells represent another step forward in providing a source of islets for the treatment of T1D with the long-term aim of eliminating the need for encapsulation or systemic immunosuppression.

### Engineering hypoimmunogenic SC-islet cells

Since GAPDH is constitutively expressed in all cells of the human islet (**Figure S1A & B**), we targeted the expression of PD-L1 and the HLA-E long-chain fusion to the GAPDH locus of hESCs and used luminescence as a reporter of cell viability. We chose to over-express PD-L1 as it was previously shown to protect SC-islet cells from xeno-rejection (Yoshihara et al., 2020), while the HLA-E long-chain fusion inhibits NK cells in a HLA-deficient context (Gornalusse et al., 2017; Mattapally et al., 2018; Riolobos et al., 2013; Wang et al., 2015). GAPDH-targeting plasmids were modified to include PD-L1 or HLA-E (**Figures S1C-E**) and used to create five hESCs lines (**Figure 1A**) - wild-type (WT), HLA-deficient (B2M^-/-^), PD-L1-expressing (PD-L1) and HLA-deficient/PD-L1-expressing (B2P) and HLA-deficient/HLA-E-expressing (BEC). All five gene-modified hESC lines successfully differentiated into SC-β cells with similar efficiency as assessed by C-peptide and Nkx6.1 expression (**Figure 1B**). Knockout of the beta 2 microglobulin (B2M) gene effectively eliminated HLA class I expression on B2M^-/-^, B2P and BEC SC-islet cells. After IFN-γ stimulation, HLA class I and PD-L1 were upregulated in WT SC-islet cells (Castro-Gutierrez et al., 2021), whereas B2P SC-islet cells constitutively over-expressed PD-L1 and lacked HLA class I expression (**Figure 1C**). A similar expression profile of HLA-E and HLA class I expression was observed on WT and BEC SC-islet cells (**Figure 1D**) (Gornalusse et al., 2017). Immunohistochemistry of magnetically-enriched SC-β cells showed localization of the HLA-E long-chain fusion to the membrane of BEC SC-β cells (**Figure 1E**) (Veres et al., 2019).

**Figure 1:**
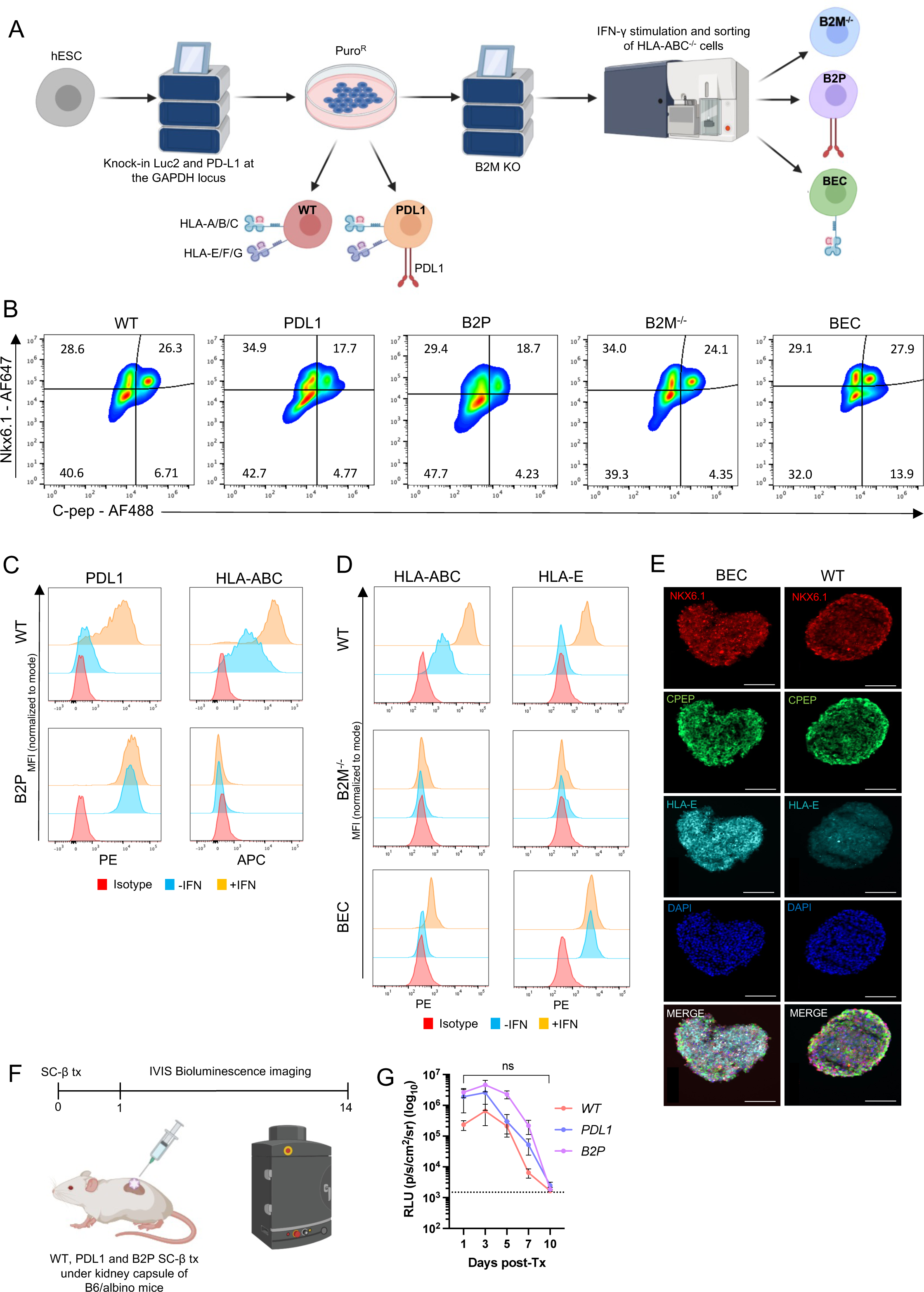
Generation of immune-evasive SC-islet cells. A. Schematic representation of the genetic-engineering strategy to generate hypoimmunogenic hESCs. B. FACS analysis of Nkx6.1/C-peptide SC-β cells derived from hypoimmunogenic hESCs (S6d14). C. PDL1 and HLA-ABC expression on SC-β cells. Data is presented as MFI (gated on C-pep^+^/Nkx6.1^+^ SC-β cells) normalized to mode. D. HLA-ABC and HLA-E expression on SC-β cells. Data is presented as MFI (gated on C-pep^+^/Nkx6.1^+^ SC-β cells) normalized to mode. E. Immunofluorescence staining of HLA-E, Nkx6.1 and C-peptide in CD49a^+^-enriched SC-β cells (S6d14). Size bars = 100μm. F. Schematic of *in vivo* SC-islet cell xeno-rejection assay. G. Quantitative analysis of xenograft rejection. Data is presented as mean ± SEM (*n* = 5/group). Dashed line represents background luminescence.

To confirm that human PD-L1 expressed on the surface of SC-islet cells binds PD-1, we assessed binding of fluorescently labelled, soluble human and mouse PD-1-Fc on WT and B2P SC-islet cells (**Figure S1F**). Both soluble human and mouse PD-1 bind membrane-bound PD-L1 on B2P SC-islet cells, whereas WT SC-islet cells do not endogenously express sufficient levels of PD-L1 to detectably bind human or mouse PD-1. We also observed a decrease in the binding affinity of soluble mouse PD-1 to human PD-L1 (Freeman et al., 2000). When WT, PD-L1 and B2P SC-islet cells were transplanted under the kidney capsule of B6/albino mice (**Figure 1F**), all gene-modified SC-islet cells were rejected within 10 days after transplantation (**Figure 1G**). These results suggest that PD-L1 over-expression is not sufficient to protect HUES8-derived SC-islet cells from xeno-rejection, and that ablation of HLA expression does not improve xenograft survival in our animal model of xeno-rejection.

### HLA-deficient SC-islet cells are resistant to allogeneic immune cell destruction in vitro

We next assessed the *in vitro* survival of gene-modified SC-islet cells in co-culture with allogeneic human immune cells (**Figure 2A**). In concordance with previous studies, NK cells and CD4^+^ and CD8^+^ T cells represented ∼8, 35 and 10% of enriched PBMCs respectively (**Figure S2A & B**) (Kleiveland, 2015). When SC-islet cells were pre-treated with IFN-γ and co-cultured with human PBMCs at multiple effector:target ratios, B2M^-/-^ and BEC SC-islet cells demonstrated significantly improved survival in comparison to WT (**Figures 2B** **& C**). HLA-E over-expression did not provide additional protective benefit. Additionally, similar survival patterns were observed when SC-islet cells were co-cultured with purified CD8^+^ and CD4^+^ T cells (**Figures S2C-F**). In all co-culture assays, there was no significant difference in the survival of all gene-modified SC-islet cells cultured with CD3/CD28 activated cells.

**Figure 2:**
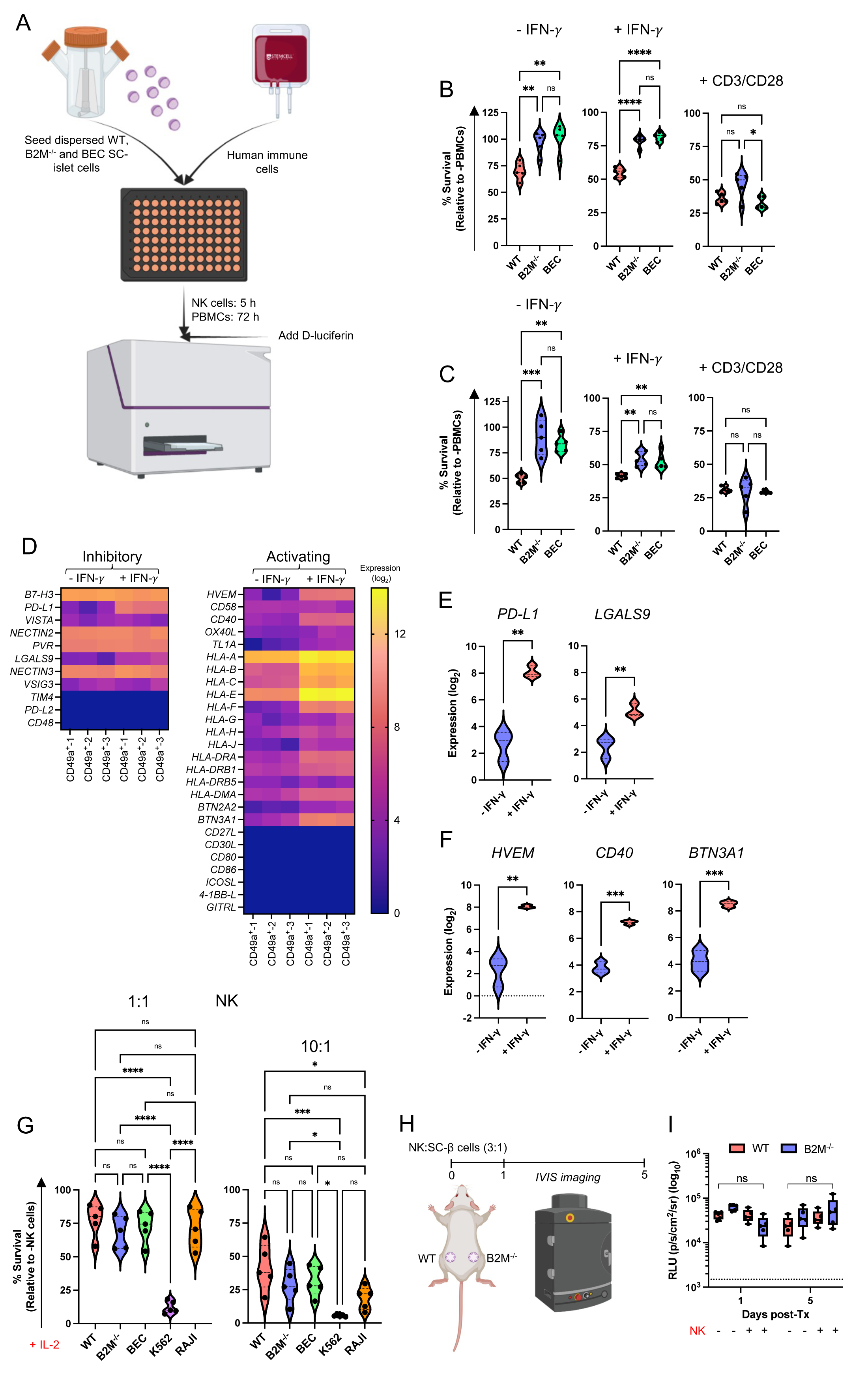
*In vitro* co-culture of hypoimmunogenic SC-islet cells with allogeneic human immune cells. A. Schematic of the *in vitro* immune:SC-islet cell co-culture assay. B. Quantification of SC-islet cell survival when co-cultured with primary human PBMCs at a 1:1 ratio. Cell survival is presented as mean ± SD (*n* = 5). C. Quantification of SC-islet cell survival when co-cultured with primary human PBMCs at a 3:1 ratio. Cell survival is presented as mean ± SD (*n* = 5). D. Heatmap of T cell ligand gene expression in IFN-γ treated CD49a^+^ SC-β cells. Ligand expression is presented as fold-change (log_2_). E. Differential expression of T cell co-inhibitory ligands in IFN-γ treated CD49a^+^ SC-β cells. Ligand expression is presented as fold-change (log_2_). F. Differential expression of T cell co-activating ligands in IFN-γ treated CD49a^+^ SC-β cells. Ligand expression is presented as fold-change (log_2_). G. Quantification of SC-β cell survival when co-cultured with primary human CD56^+^ NK cells. Cell survival is presented as mean ± SD (*n* = 5). H. Schematic of *in vivo* NK cell cytotoxicity assay. I. Quantitative analysis of *in vivo* NK cell assay. Data is presented as mean ± SEM. Dashed line represents background luminescence.

Since expression of T cell co-activating and co-inhibitory ligands dictates T cell function and is regulated by various stimuli including IFN-γ stimulation (Chen and Flies, 2013), we performed bulk RNA sequencing on WT CD49a^+^ SC-β cells and assessed T cell ligand expression after IFN-γ stimulation (**Figure 2D**). As expected, IFN-γ stimulation upregulated the T cell co-inhibitory ligands *PD-L1* and *LGALS9* (galectin 9) in SC-β cells (**Figure 2E**) (Garcia-Diaz et al., 2017; Imaizumi et al., 2002). Conversely, while we did not detect transcripts for many T cell co-activating ligands other than HLA genes, we found that IFN-γ stimulation upregulated members of the TNF receptor superfamily *HVEM* (TNF Receptor Superfamily Member 14) and *CD40* (Tumor Necrosis Factor Receptor Superfamily Member 5) in SC-β cells (**Figure 2F**) (Benci et al., 2016; Wagner et al., 2002). Interestingly, *BNT3A1* (butyrophilin subfamily 3 member A1), which is an MHC-associated gene that regulates T cell activation and proliferation (Rigau et al., 2020), was also upregulated. These results suggest that in the absence of classical HLA-TCR signaling due to HLA knockout, other co-stimulatory and co-inhibitory T cell ligands are expressed in SC-β cells that may influence T cell function.

### HLA-deficient SC-islet cells are resistant to allogeneic NK cell destruction in vitro and in vivo

We next assessed the effect of HLA deletion and HLA-E over-expression on NK cell function against SC-islet cells. When co-cultured with NK92mi cells, BEC SC-islet cells showed no significant difference in survival in comparison to WT SC-islet cells, whereas HLA-deficient SC-islet cells were susceptible to NK92mi cytotoxicity (**Figure S3A**) (Gornalusse et al., 2017). The strong protective effect of HLA-E against NK92mi cells is likely due to the high percentage (∼96.1%) of NKG2A^+^/NKG2C^-^ cells (**Figure S3B)**, which biases NK cell inhibition.

While NK cell lines are useful tools for assessing NK cell cytotoxicity, they do not accurately recapitulate primary NK cell receptor expression and function. Previous studies have shown that in the absence of IL-2 activation, human NK cells do not destroy HLA-deficient endothelial cells and platelets (Deuse et al., 2021; Suzuki et al., 2020). Thus, prior to co-culture with gene-modified SC-islet cells, we pre-activated human NK cells with IL-2 for 5 days as previously described (Deuse et al., 2021). Enriched NK cells consisted of ∼80% CD56^dim^ and ∼5% CD56^high^ NK cells (**Figure S3C & D**). Surprisingly, despite IL-2 pre-activation, we observed no significant difference in survival between gene-modified SC-islet cells at multiple effector:target ratios (**Figure 2G**). Since HLA-E over-expression does not provide any additional protective benefit to HLA-deficient SC-islet cells against NK cell cytotoxicity, we chose to interogate WT and B2M^-/-^ SC-islet cells moving forward. We assessed the survival of WT and B2M^-/-^ SC-islet cells after transplantation with IL-2 pre-activated NK cells in Scid/beige mice (**Figure 2H**). Again, there was no significant difference in WT and B2M^-/-^ SC-islet cell survival *in vivo* (**Figure 2I** **& S3E**). Taken together, these results suggest that HLA-deficient SC-islet cells are intrinsically resistant to NK cell cytotoxicity and that their survival *in vitro* is recapitulated *in vivo*.

### Exclusively, SC-β cells intrinsically possess and sustain an NK cell evasive ligand profile after inflammatory stimulus

Since NK cell function is dictated by a balance of inhibitory and activating signals, we compared the ligand profile of CD49a^+^ SC-β cells to stem cell-derived endothelial (SC-endo) cells. We chose endothelial cells as a comparative cell type because it has previously been shown that HLA-deficient endothelial cells are susceptible to pre-activated NK cells (Deuse et al., 2021). Endothelial cell differentiation resulted in >95% CD31^+^ SC-Endo cells derived from both WT and B2M^-/-^ hESCs (**Figures S3F & G**). Bulk RNA sequencing and transcript analysis of NK cell ligands in SC-β and SC-endo cells confirmed that SC-endo cells possess an activating NK cell ligand profile, characterized by expression of *MICA*, *MICB*, *ULBP1*, *ULBP2* and *RAET1G*, whereas transcripts for these genes could not be detected in SC-β cells (**Figures 3A** **& B**). Importantly, we found that the expression of most NK cell ligands is not IFN-γ regulated, except for the HLA molecules. Thus, within an HLA-deficient context, the expression of non-HLA NK cell ligands is stable following an inflammatory stimulus. We confirmed the bulk RNA sequencing analysis of NK cell ligand expression by flow cytometry, which showed that MIC and ULBP proteins were either absent or less expressed on SC-β cells (**Figure 3C** **& D**). Taken together these data suggest that SC-β cells intrinsically possess and sustain a diminished NK cell activating ligand profile after inflammatory stimulus, and that this ligand phenotype may explain their resistance to IL-2 pre-activated NK cell cytotoxicity.

**Figure 3:**
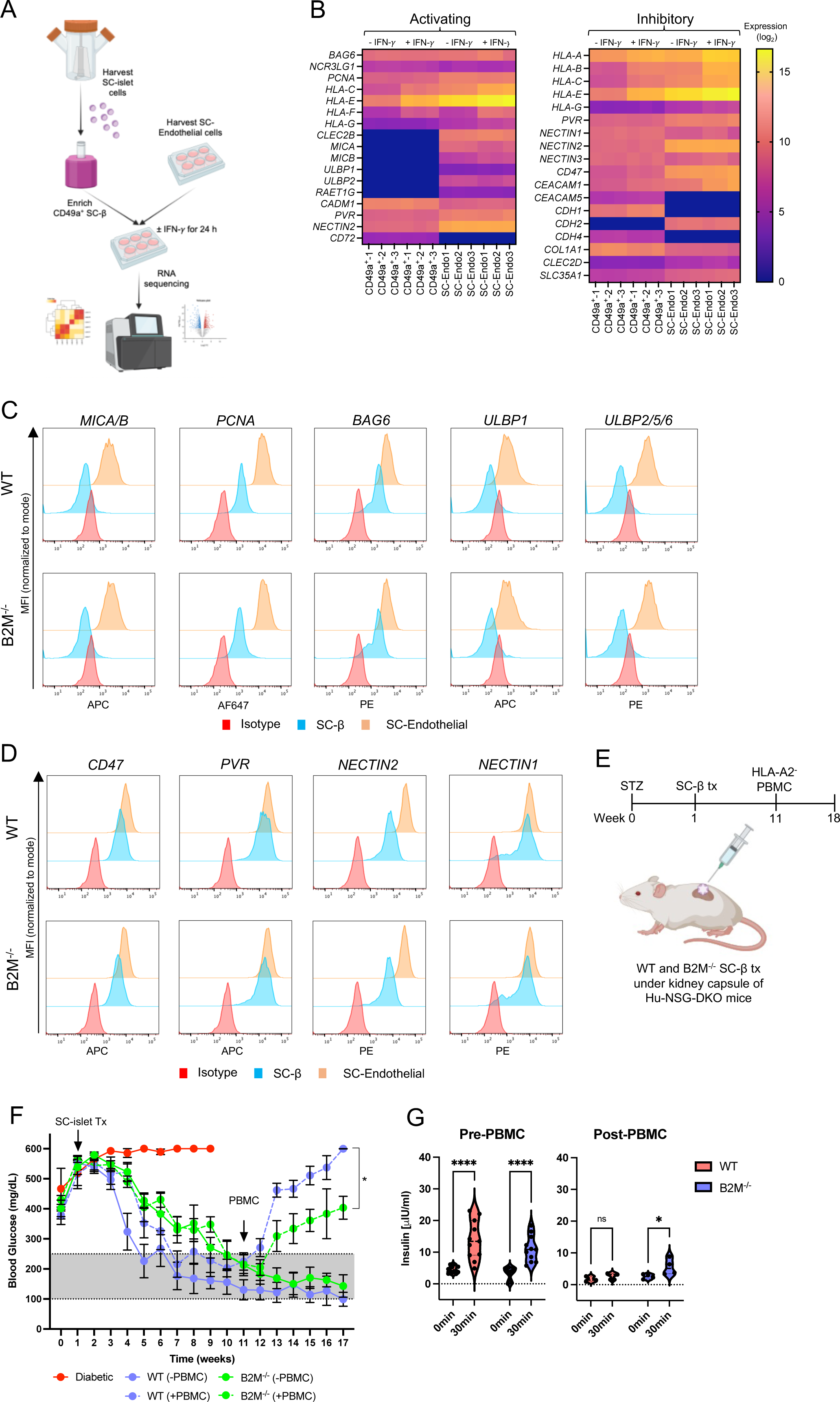
Characterization of SC-islet cell-specific NK cell ligand profile. A. Schematic of SC-β and SC-Endothelial cell bulk RNA sequencing workflow. B. Heatmap of NK cell ligand expression in SC-β and SC-Endothelial cells (-IFN-γ vs +IFN-γ). Ligand expression is presented as fold-change (log_2_). C. NK cell activating ligand expression on SC-β and SC-Endothelial cells. Data is presented as MFI normalized to mode and is representative of three independent experiments. D. NK cell inhibitory ligand expression on SC-β and SC-Endothelial cells. Data is presented as MFI normalized to mode and is representative of three independent experiments. E. Schematic of SC-islet cell transplantation in diabetic humanized NSG-DKO mice. F. Non-fasting blood glucose concentrations after SC-islet transplantation and PBMC injection. Data is presented as mean ± SD (*n* = 12/group). G. In vivo GSIS pre- and post-PBMC injection. Plasma insulin concentration was measured t=0 and 30 min after glucose injection. Data is represented as mean ± SD (*n* = 5/group).

### HLA-deficient SC-islet cells reverse diabetes in humanized mice and demonstrate delayed graft rejection

Having demonstrated that HLA-deficient SC-islet cells resist PBMC cytotoxicity *in vitro* and NK cell cytotoxicity *in vitro* and *in vivo*, we assessed the survival of HLA-deficient SC-islet cells in diabetic, NSG-DKO mice prior to PBMC injection (**Figure 3E**). By 10 weeks post-transplantation there was no significant difference in blood glucose levels of mice transplanted with WT and B2M^-/-^ SC-islet cells (**Figure 3F**). Once blood glucose levels had been normalized by the SC-islet transplants, we injected HLA-A2^-^ (mis-matched) PBMCs and assessed allograft rejection by monitoring blood glucose levels. WT SC-islet cells were destroyed within 2 weeks, whereas the rejection of B2M^-/-^ SC-islet cells was delayed. At 7 weeks post-PBMC injection, *in vivo* GSIS assay showed that animals transplanted with WT SC-islet cells lost *in vivo* graft function, while some graft function was observed in animals transplanted with B2M^-/-^ SC-islet cells (**Figure 3G**). These results suggest that B2M^-/-^ SC-islet cells reverse diabetes and demonstrate delayed allograft rejection, however it is likely that they would ultimately be completely rejected.

### Cytokine-secreting SC-β cells induce a local tolerogenic microenvironment that protects against xeno-rejection

As a result of the varying success of immune-evasive engineering to protect SC-islet cells, we pursued a complementary strategy by engineering SC-islet cells to secrete the cytokines IL-2 mutein, TGF-β and IL-10 as vehicles of localized immune (**Figures 4A**). The IL-2 mutein (N88D) possesses reduced affinity for the IL-2Rβγ receptor, resulting in the production of a Treg-selective molecule that preferentially expands T_regs_ while having a minimal effect on CD4^+^ and CD8^+^ memory T cells (Khoryati et al., 2020; Peterson et al., 2018). Since the immune-suppressive phenotype of T_regs_ requires IL-10 and TGF-β, we included these two cytokines in our immune-tolerizing construct (Horwitz et al., 2008).

**Figure 4:**
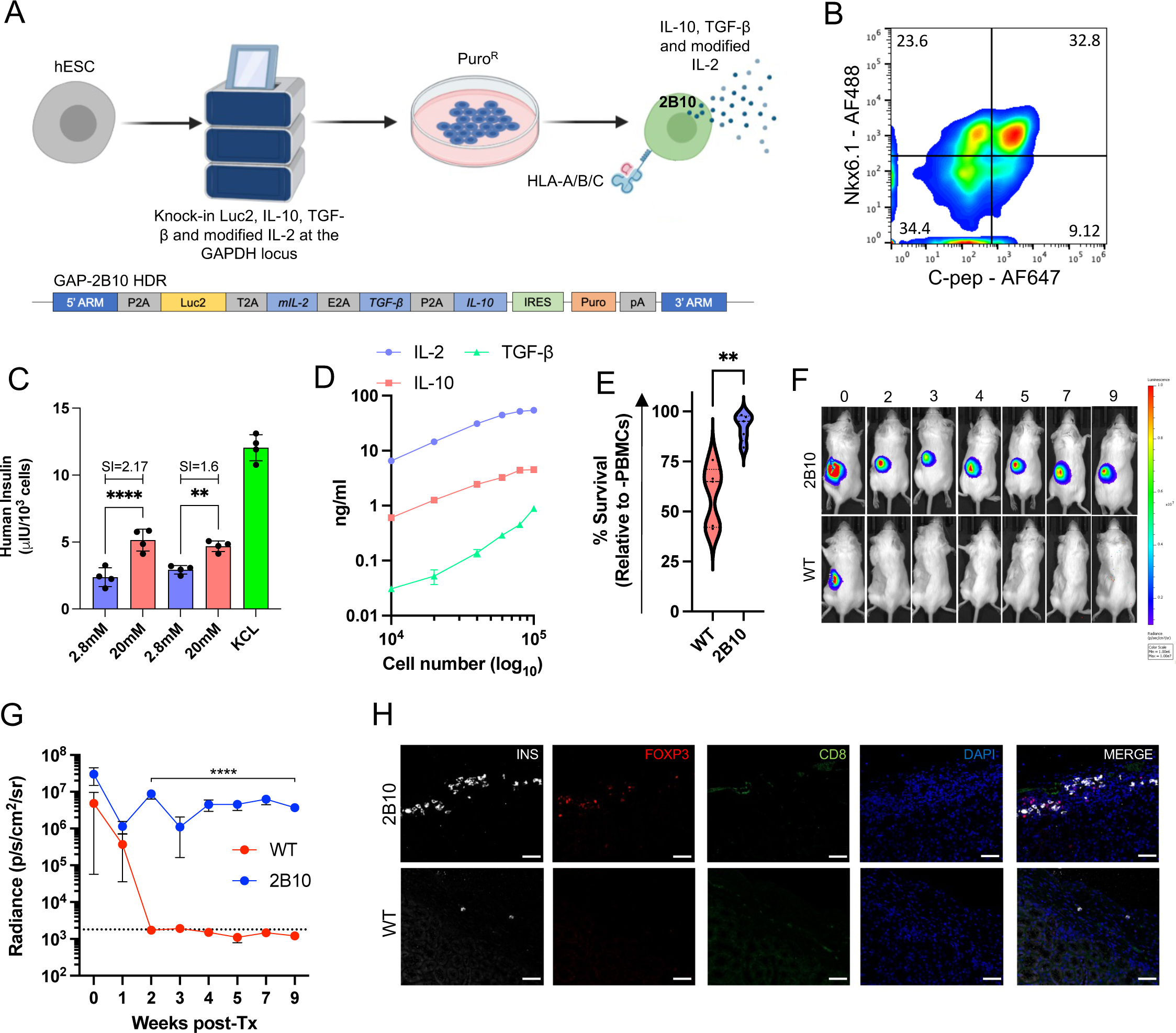
Immune-tolerizing SC-islet cells survive xeno-rejection. A. Schematic representation of the genetic-engineering strategy to generate immune-modulatory hESCs and of the GAP-2B10 HDR plasmid. B. FACS analysis of Nkx6.1/C-peptide SC-β cells derived from immune-modulatory hESCs (S6d8). C. In vitro GSIS of 2B10 SC-islet cells. Data is represented as mean ± SD (*n* = 4). D. Quantification of cytokine secretion from 2B10 SC-islet cells. Data is represented as mean ± SD (*n* = 2). E. Quantification of SC-islet cell survival when co-cultured with primary human PBMCs at a 1:1 ratio. Cell survival is presented as mean ± SD (*n* = 5) F. In vivo bioluminescence imaging of B6/albino mice transplanted with 2B10 SC-islet cells (*n* = 3/group). G. Quantitative analysis of *in vivo* graft survival in B6/albino mice. Data is presented as mean ± SD (n = 3/group). H. Immunostaining of 2B10 and WT SC-islet grafts showing presence of INS^+^ cells and Treg recruitment at 5 weeks post transplantation. Size bars = 100μm.

Differentiation of this genetically-modified cell line, called 2B10, resulted in ∼30% Nkx6.1^+^/C-peptide^+^ SC-β cells (**Figure 4B**) with physiological function as assessed *in vitro* by GSIS (**Figure 4C**). We also confirmed that 2B10 SC-islet cells secrete modified IL-2, TGF-β and IL-10, and that the quantity was cell-concentration dependent (**Figure 4D**). When WT and 2B10 SC-islet cells were co-cultured with human PBMCs in *vitro*, 2B10 SC-islet cells showed significantly improved survival (**Figure 4E**). Notably, 2B10 SC-islet cells transplanted under the kidney capsule of B6/albino mice survived up to 9 weeks post-transplantation, while WT SC-islet cell were destroyed within 2 weeks (**Figures 4F** **& G**). Furthermore, we found that WT grafts contained little to no remaining SC-islet cells, while 2B10 grafts contained surviving insulin-producing cells and T_regs_ localized within the graft (**Figure 4H**). Collectively, these data suggest that SC-islet cells can be engineered to co-secrete immune-modulatory cytokines that induce a tolerogenic local microenvironment characterized by Treg cell infiltration that sustains xenograft survival.

## Discussion

The utility of an immune-evasive/tolerogenic islet cell replacement therapy relies on the ability to maintain transgene expression throughout cell differentiation and after transplantation. Thus, to analyze the effect of various gene-editing strategies for the immune protection of SC-islet cells, we engineered SC-islet cells to constitutively express tolerogenic molecules including PD-L1 (Castro-Gutierrez et al., 2021; Yoshihara et al., 2020) and the HLA-E long-chain fusion (Gornalusse et al., 2017) from the *GAPDH* locus, both in a HLA-competent and HLA-deficient background. We report the lack of a xeno-protective effect of PD-L1 over-expression in SC-islet cells. This may be explained by species-specific differences in PD-L1/PD-1 binding (Viricel et al., 2015), and the fact that human and mouse PD-1 share only 60% homology at the amino acid level (Finger et al., 1997). In fact, we observed decrease binding of soluble mouse PD-1 to human PD-L1 expressed on SC-islet cells. While these results suggest that over-expression of human PD-L1 is not sufficient to overcome xeno-rejection, PD-L1 over-expression may still be useful in an allogeneic setting.

In concordance with previous studies, HLA-deficient SC-islet cells were resistant to PBMC cytotoxicity *in vitro* (Han et al., 2019; Leite et al., 2022). We also found that SC-β cells modulate their T cell ligand profile in response to partial inflammatory stimulus. While we and others have attempted to exploit the PD-L1/PD-1 T cell signaling axis to engineer immune-evasive SC-islet cells (Yoshihara et al., 2020), our transcript analysis of T cell ligands in SC-β cells suggests that exploiting the LGALS9/TIM-3 signaling axis may also be of interest. In fact, like *PD-L1*, *LGALS9* is frequently upregulated in cancer cells where it contributes to tumor progression by inhibition of T cell function (Heusschen et al., 2013; Yang et al., 2021). Additionally, we identified the T cell activating ligands *HVEM*, *CD40* and *BTN3A1* as potential targets to knock-out in SC-islet cells to further influence T cell function.

We also showed that HLA-deficient SC-islet cells are resistant to pre-activated NK cell cytotoxicity both *in vitro* and *in vivo*, and that over-expression of the HLA-E long-chain fusion does not provide any additional protective benefit. This is due to the SC-islet cell-specific lack of NK cell activating ligands such as the MIC and ULBP proteins, which are expressed in SC-Endothelial cells, and may explain why SC-Endothelial cells are susceptible to pre-activated NK cell cytotoxicity while SC-islet cells are resistant (Deuse et al., 2021). However, since many immunodeficient mouse models lack critical components for NK cell survival and function such as SIRPα and IL-15 (Herndler-Brandstetter et al., 2017), they do not support long-term engraftment of human NK cells and we cannot exclude the possibility of HLA-deficient SC-islet cell destruction in a mouse model that better supports NK cell engraftment.

Furthermore, while transplantation of HLA-deficient SC-islet cells was able to normalize blood glucose levels in diabetic mice, after PBMC injection the cells were eventually rejected, albeit with delayed rejection kinetics. This could be explained by the presence of other immune cell subsets such as macrophage, monocytes and dendritic cells that may play a role in indirect allograft rejection (Oberbarnscheidt et al., 2014; Wyburn et al., 2005; Zhuang et al., 2016). Ultimately, this highlights the limitations of exclusively using *in vitro* immune cell co-culture assays to assess the effect of genetic modification on the protection SC-islet cells from immune destruction (Castro-Gutierrez et al., 2021; Leite et al., 2022).

Finally, we show for the first time that SC-islet cells engineered to secrete modified IL-2, TGF-β and IL-10 are protected against xeno-rejection. This finding has several important implications. First, since β cells are professional secretory cells with a significant translatory demand, it demonstrates that SC-islet cells can be co-opted to secrete other proteins while maintaining their designed function (Lim et al., 2020).

Second, the intrinsic ligand profile of the desired cell type should be considered when determining the set of genetic modifications required to generate immune-evasive cells, as some cell types may not require extensive genetic manipulation. Since the safety of immune-evasive/immune-tolerizing cell therapies is critical to their clinical translation, ideally the genetically engineered product should be generated with the least number of genetic perturbations to limit off-target events and chromosomal instability. Third, our results provide validation for the use of immune-tolerizing approaches (either alone or in conjunction with immune-evasion) as a method to protect SC-islet cells from the immune system. A variation of this approach may be to include some, but not all cells, in the transplant that secrete these tolerizing molecules. Overall, this approach may eliminate the need for encapsulation or immunosuppression, a long-standing goal of the islet transplantation field.

## Limitations

Although HLA-deficient SC-islet cells possessed improved survival in *in vitro* PBMC co-culture assays (**Figures 2B and C**), we found that these results did not translate *in vivo*. Choosing an appropriate humanized mouse model is essential to evaluate immune-evasion/immune-tolerizing genetic engineering strategies. Many PBMC humanized mouse models such as that used in this study do not fully recapitulate human allograft rejection since they bias CD3^+^ cell engraftment and lack other immune cell subsets. Thus, while HLA-deficient SC-islets demonstrated delayed rejection in the PBMC humanized mouse model (**Figure 3F**), we cannot exclude the possibility that in CD34^+^ or BLT mice (humanized mouse models that better recapitulate human immune components), that these cells would be rejected earlier. For these reasons, validating the protective effect of immune-evasion/tolerizing genetic-engineering strategies should ideally be conducted in humanized mouse models that more accurately reflect allograft rejection. While xeno-rejection does not mimic allo-rejection, we ultimately chose to use the xeno-rejection model to evaluate our immune-tolerizing genetic-engineering strategy as it is a stronger model of immune rejection and eliminates the immune cell bias of humanized mouse models.

By using such a strong model of graft rejection, additional immune barriers are introduced that require complex genetic-engineering approaches as a solution. This means combining gene knock-outs and knock-ins to generate the desired immune-evasive/tolerizing cell product. Knock-in of multiple genes at a specific locus in the genome requires large homology directed repair templates, which is associated with poor integration efficiency and recovery of genetically unstable clones. Exploring the possibility of introducing tolerogenic molecules at multiple constitutively expressed loci could reduce the size of HDR templates and result in the recovery of genetically stable homozygous knock-in clones. Striking a balance between HDR length and the number of editing events may permit the introduction of more foreign genetic material into the genome without destabilizing effects.

Furthermore, since immune-tolerizing SC-islets secrete cytokines from all cells within the islet, there is a risk that the concentration of constitutively secreted cytokines may result in chronic immunosuppression. Thus, it is important to evaluate the immune status of mice receiving immune-tolerizing SC-islet cells as chronic immunosuppression should be avoided. However, if transplantation of immune-tolerizing SC-islets does result in chronic immunosuppression, the ability to enrich for specific endocrine cell populations may allow us to adjust the dose of cytokine-secretion by generating designer islets composed of cytokine-secreting and non-cytokine secreting endocrine cells such that localized graft tolerance is achieved.

## Methods

### Mice

Male B6/albino, Scid/beige and NSG-MHC Class I/II KO mice (6-8 weeks of age) were purchased from Jackson Labs. Mice were housed in specific pathogen-free conditions at Harvard University. All animal research was conducted under Harvard IACUC approval.

### Cell Culture

Human ES maintenance and differentiation was carried out as previously described (Millman et al., 2016; Pagliuca et al., 2014). SC-islet cell differentiations were initiated 72 h after initial passage by aspirating mTeSR1 and replenished with stage and day-specific media supplemented with the appropriate small molecules or growth factors as previously described (Millman et al., 2016; Pagliuca et al., 2014; Veres et al., 2019). All cell lines were routinely tested for mycoplasma and were mycoplasma-free. All experiments involving human cells were approved by the Harvard University IRB and ESCRO committees.

### Generation of immune-evasive hESCs

The existing GAPluc (WT) hESC line served as starting material for the generation of HLA-deficient hESCs (Gerace et al., 2021). To generate HLA-deficient hESCs, 1x10^6^ WT hESCs were nucleofected using the 4D-Nucleofector (Lonza) with RNP complexed with 120 pmol B2M gRNA (5’-GCTACTCTCTCTTTCTGGCC’3)(Mandal et al., 2014) (IDT) and 104 pmol Alt-R® S.p. HiFi Cas9 Nuclease V3 (IDT) according to the manufacturer’s instructions. Nucleofected cells were resuspended in mTesR1 (STEMCELL Technologies, 85850) + 10 µM Y27632 (DNSK International, DNSK-KI-15-02) and plated in a matrigel-coated tissue culture plate. After 48 h cells were treated with 10ng/ml IFN-γ (R&D, 285-IF-100) for 24 h and then stained with APC anti-human HLA-ABC (W6/32, 1:100) (Biolegend, 311409). HLA-ABC^-/-^ cells were sorted on a FACS Aria II (BD Biosciences) and plated in a matrigel-coated tissue culture plate containing mTesR1 + CloneR (STEMCELL Technologies, 05888), with single colonies picked for expansion. The resulting HLA-deficient line was named B2M^-/-^.

To generate GAP-PD and GAP-BEC hESCs, human PD-L1 and the peptide::B2M::HLA-E fusion sequences were synthesized as gBlocks (Genscript) and cloned into our existing GAPluc targeting plasmid downstream of the *Luc2* gene. The sequence of the covalently attached peptide (VMAPRTLLL) in the peptide::B2M::HLA-E fusion is derived from the HLA-Cw7 molecule (Kaiser et al., 2005). hESCs were nucleofected with the GAP-PD or GAP-BEC targeting plasmids and GAPDH-targeting RNP as previously described (Gerace et al., 2021). For GAP-B2P and GAP-BEC lines, a polyclonal population of puromycin-selected cells was subsequently nucleofected with B2M-targeting RNP, and single-colonies were picked for expansion after FACS sorting as described above. The gating strategy for sorting HLA-ABC^-/-^ hESCs is described in **Figure S1G**.

### Flow Cytometry

Differentiated WT, B2M^-/-^, BEC, PD-L1 and B2P SC-islet cells were both treated and untreated with 10ng/ml IFN-γ for 24 h prior to staining. Cells were dissociated with Accutase (STEMCELL Technologies, 07920), washed twice with PBS + 0.1% BSA (Gibco, A10008-01) and blocked for 30 min on ice with PBS + 5% donkey serum (Jackson Labs; 100181-234). Cells were then stained for 30 min on ice in blocking buffer with PE or APC anti-human HLA-ABC (Biolegend, 311405), PE anti-human HLA-E (Biolegend, 342603) and PE anti-human PD-L1 (Biolegend, 393607). Cells were then washed three times and fixed in 4% PFA (EMS, 15710) for 15 min at 4 °C. Fixed cells were then incubated in blocking buffer with rat anti-human C-peptide (DHSB; GN-ID4) and mouse anti-human Nkx6.1 (DHSB; F55A12) (overnight at 4 °C), washed three times with blocking buffer, incubated with goat anti-rat 647 (Life Technologies, A-21247; 1:300) and goat anti-mouse 405 (Life Technologies, A-31553; 1:300) in blocking solution (1 h at room temperature), washed three times and resuspended in PBS + 0.1% BSA. Samples were captured on the LSR II (BD) flow cytometer and analyzed using FlowJo 10.7.1 (BD). All antibodies were used at 1:100 unless otherwise stated. The gating strategy for identifying SC-β cells is described in **Figure S1H**. PE mouse IgG2a (Biolegend, 400213) and PE mouse IgG1 (Biolegend, 400111) served as isotype controls.

### Magnetic enrichment using CD49a and reaggregation

WT and BEC SC-β cells were magnetically-enriched from SC-islet clusters as previously described (Veres et al., 2019). Enriched cells were resuspended in S6 medium and plated at 5x10^3^ cells/well in low-attachment 96-well v-bottom tissue-culture plates (Thermo Scientific, 277143), centrifuged at 300 g for 1 min and incubated at 37 °C for 4-7 days. CD49a enriched SC-β cell clusters were fed fresh S6 medium every 2 days.

### SC-β cell immunohistochemistry

CD49a enriched SC-β cell clusters were fixed in 4% PFA for 1 h at room temperature, washed and frozen in OCT (Tissue-Tek, 4583) and sectioned to 14 µm. Before staining, paraffin-embedded samples were treated with Histo-Clear (EMS, 64110-01) to remove the paraffin. For staining, slides were incubated in blocking buffer (PBS + 0.1% Triton-X + 5% donkey serum) for 1 h at room temperature, incubated in PBS + 5% donkey serum containing rat anti-human C-peptide (1:300), mouse anti-human Nkx6.1 (1:100) and rabbit anti-human HLA-E (Sigma-Aldrich, HPA031454, 1:100) for 1 h at room temperature, washed three times, incubated in goat anti-mouse 594 (Life Technologies, A-11032; 1:500), goat anti-rat 488 (Life Technologies, A-11006; 1:500) and goat anti-rabbit 647 (Life technologies, A-21244; 1:500) for 2 h at room temperature, washed, mounted in Vectashield with DAPI (Vector Laboratories; H-1200), covered with coverslips and sealed with clear nail polish. Representative regions were imaged using Zeiss.Z2 with Apotome microscope.

### Immune cell isolation

PBMCs, CD8 and CD4 T cells, and NK cells were isolated from apheresis leukoreduction collars (*n* = 5 donors) obtained from Brigham and Women’s Hospital in compliance with our IRB approval. PBMCs were isolated by density gradient centrifugation with lymphoprep (STEMCELL Technologies, 07801) SepMate™-50 (IVD) tubes (STEMCELL Technologies, 85450) according to the manufacturer’s instructions. CD4, CD8 and NK cells were isolated by density gradient centrifugation with lymphoprep and SepMate™-50 (IVD) tubes following the addition of RosetteSep Human CD4+ T Cell Enrichment Cocktail (STEMCELL Technologies, 15062), CD8+ T Cell Enrichment Cocktail (STEMCELL Technologies, 15063) and NK Cell Enrichment Cocktail (STEMCELL Technologies, 15065) respectively. Enriched immune cells were cryopreserved in CryoStor CS10 (STEMCELL Technologies, 07930) at a concentration of 10M cells/vial.

Flow cytometric analysis of enriched immune cell subpopulations was performed by staining cells in blocking buffer containing APC anti-human CD3 (Biolegend, 300311), PE anti-human CD4 (Biolegend, 357403), Pacific Blue anti-human CD8 (Biolegend, 344717) and Pacific Blue anti-human CD56 (Biolegend, 362519). Isotype controls were Pacific Blue mouse IgG1 (Biolegend, 400131), APC mouse IgG2a (Biolegend, 400221) and PE rat IgG2b (Biolegend, 400607). All antibodies were used at 1:100 or unless otherwise stated. Representative gating strategies are demonstrated in **Figures S2G-I**.

### In vitro T cell cytotoxicity assays

WT, B2M^-/-^ and BEC SC-islet cells were seeded in S3 medium at 5 x 10^4^ cells/well of a matrigel-coated 96-well black flat-bottom plate (Corning, 3916) in the presence or absence of IFN-γ (10 ng/ml). After 24 h, the medium was switched to T cell medium (ImmunoCult™-XF T Cell Expansion Medium (STEMCELL Technologies, 10981) + 100U/ml rhIL-2) for immune cell co-culture. Primary human PBMCs, CD4 or CD8 cells (*n* = 5 donors) cultured in T cell medium were added to SC-islet cells at 1:1 and 3:1 effector:target ratios with and without the addition of ImmunoCult™ Human CD3/CD28 T Cell Activator (STEMCELL Technologies, 10991). All T cell cytotoxicity assays were co-cultured for 72 h, after which luminescence was measured following the addition of 150 µg/ml D-luciferin (Gold Biotechnology, LUCK-2G) on the CLARIOstar microplate reader (BMG LABTECH). SC-islet cell survival was calculated as a percentage relative to luminescence in the absence of T cells (test SC-islet cell luminescence/no T cell SC-islet cell luminescence x 100).

### In vitro NK cell cytotoxicity assays

WT, B2M^-/-^ and BEC SC-islet cells were seeded in NK cell medium (NK MACS medium (Milteny Biotec, 130-114-429) + 5% Human AB serum (Valley Biomed, HP1022HI) + 5% HyClone FBS (GE Healthcare, SH30070.03) + 0.5 ng/ml rhIL-2 (Peprotech, 200-02)) at 2 x 10^4^ cells/well of a 96-well black round-bottom ultra-low attachment plate (Corning, 4591). Primary NK cells (*n* = 5 donors) cultured in NK cell media for 5 days were added to SC-islet cells at 1:1 and 10:1 effector:target ratios. K562-Luc2 (Biocytogen, BCG-PS-015-luc) and Raji-GFP-Luc2 (Biocytogen, BCG-PS-087-luc) cells were used as positive and negative controls respectively. All NK cell cytotoxicity assays were co-cultured for 5 h, after which luminescence and SC-islet cell survival was measured as described above.

### In vivo NK cell cytotoxicity

2.5 x 10^5^ WT and B2M^-/-^ SC-islet cell clusters were resuspended either alone or with 7.5 x 10^5^ primary human NK cells (pre-treated with 0.5 ng/ml rhIL-2 for 12 h) in phenol-free Matrigel (Corning, 356237) and transplanted subcutaneously in Scid/beige mice (*n* = 5). Bioluminescence was measured 10 min after i.p injection of 10 μL/g D-luciferin (15mg/ml) on days 1 and 5 following cell transplantation on the IVIS Spectrum (PerkinElmer, 124262) as previously described (Gerace et al., 2021).

### Endothelial cell differentiation

WT and B2M^-/-^ hESCs were seeded in mTesR1 + 10 μM Y27632 at a concentration of 7.5 x 10^4^ cells/well of a Matrigel-coated 6-well tissue-culture plate (Corning, 3516). Thereafter, cells were differentiated into endothelial cells using STEMdiff™ Endothelial Differentiation Kit (STEMCELL Technologies, 08005) according to the manufacturer’s instructions. Differentiation efficiency was quantified by FACS analysis after staining endothelial cells with Pacific Blue anti-human CD31 (Biologend, 303113). Pacific Blue mouse IgG1 (Biolegend, 400131) was used as an isotype control. All antibodies were used at 1:100 unless otherwise stated.

### Bulk RNA sequencing

Magnetically-enriched CD49a^+^ SC-β cells and SC-Endothelial cells were seeded in 6-well plates at 1 x 10^6^ cells/well (in triplicate) and treated with and without 10ng/ml IFN-γ for 24 h. Cells were then harvested using Accutase and the pellets snap-frozen on dry-ice. RNA extractions, library preparations and sequencing reactions were conducted at GENEWIZ, LLC. (South Plainfield, NJ, USA). Total RNA was extracted from cell pellet samples using RNeasy Plus Mini Kit (Qiagen, Germantown, MD, USA) and quantified using Qubit Fluorometer (Life Technologies, Carlsbad, CA, USA). SMART-Seq v4 Ultra Low Input Kit for Sequencing was used for full-length cDNA synthesis and amplification (Clontech, Mountain View, CA), and Illumina Nextera XT library was used for sequencing library preparation.

Multiplexed sequencing libraries were loaded on the Illumina HiSeq instrument and sequenced using a 2x150 Paired End (PE) configuration. Image analysis and base calling were conducted by the HiSeq Control Software (HCS). Raw sequence data (.bcl files) generated from Illumina HiSeq was converted into FASTQ files and de-multiplexed using Illumina’s bcl2fastq 2.17 software. One mismatch was allowed for index sequence identification.

After investigating the quality of the raw data, sequence reads were trimmed to remove possible adapter sequences and nucleotides with poor quality using Trimmomatic v.0.36. The trimmed reads were mapped to the Homo sapiens reference genome available on ENSEMBL using the STAR aligner v.2.5.2b. The STAR aligner is a splice aligner that detects splice junctions and incorporates them to help align the entire read sequences. BAM files were generated as a result of this step. Unique gene hit counts were calculated by using feature Counts from the Subread package v.1.5.2. Only unique reads that fell within exon regions were counted.

After extraction of gene hit counts, the gene hit counts table was used for downstream differential expression analysis. Using DESeq2, a comparison of gene expression between the groups of samples was performed. The Wald test was used to generate p-values and Log_2_ fold changes. Genes with adjusted p-values < 0.05 and absolute log_2_ fold changes > 1 were called as differentially expressed genes for each comparison. A gene ontology analysis was performed on the statistically significant set of genes by implementing the software g:Profiler (Raudvere et al., 2019). *The gene accession number for the unique RNAseq dataset generated in this study is GSE200021. The reviewer token for private access to this dataset is “gpkjywayfxmhjgd”*.

### NK cell ligand analysis

WT and B2M^-/-^ SC-islet cells and SC-endothelial cells were washed twice with washing buffer and blocked for 30 min on ice with blocking buffer. Cells were then stained for 30 min on ice in blocking buffer with APC anti-human CD47 (Biolegend, 323123), APC anti-human CD324 (E-Cadherin) (Biolegend, 324107), PE anti-human CD325 (N-Cadherin) (Biolegend, 350806), PE anti-human CD112 (Nectin-2) (Biolegend, 337409), APC anti-human CD155 (PVR) (Biolegend, 337617), PE anti-human CD111 (Nectin-1) (Biolegend, 340404), APC anti-human MICA/MICB (Biolegend, 320907), AF647 anti-human PCNA (Biolegend, 307912), PE anti-human BAG6 (Abcam, ab210838), APC anti-human CD48 (Biolegend, 336713), PE anti-human CD70 (Biolegend, 355103), mouse anti-human CD113 (Nectin-3) (Millipore-Sigma, MABT63), APC anti-human ULBP1 (R&D, FAB1380A), PE anti-human ULBP2/5/6 (R&D, FAB1298P), APC anti-human CLEC2D (R&D, FAB3480A) and PE anti-human CD72 (Biolegend, 316207). PE Mouse IgG1 (Biolegend, 400111), APC mouse IgG1 (Biolegend, 400121), APC mouse IgG2a (Biolegend, 400221), PE Mouse IgG2a (Biolegend, 400213), APC rat IgG1 (Biolegend, 401903), PE rabbit IgG (Cell Signaling technology, 5742S) and goat anti-mouse 488 (Thermofisher, A-11001, 1:300) served as isotype controls. All antibodies were used at 1:100 unless otherwise stated.

### Transplantation of HLA-deficient SC-islet cells

Diabetes was induced in NSG-MHC Class I/II KO mice by multiple low-dose (40 mg/kg) streptozotocin (STZ) i.p. injection as previously described (Furman, 2021). Once animals reached a blood glucose of >500 mg/dL, 5 x 10^6^ WT or B2M^-/-^ SC-islet cells were transplanted under the kidney capsule (*n* = 10) as previously described (Millman et al., 2016; Pagliuca et al., 2014). Blood glucose and body weight was measured twice a week after transplantation.

Ten weeks after transplantation (before PBMC injection) and seven weeks after PBMC injection the function of transplanted cells was assessed by performing *in vivo* peritoneal glucose-stimulated insulin secretion (IPGTT) as previously described (Millman et al., 2016; Pagliuca et al., 2014). Insulin secretion was quantified using the Human Ultrasensitive Insulin ELISA (ALPCO Diagnostics; 80-INSHUU-E01.1.)

### Generation of immune-tolerizing SC-islet cells

The cDNAs of human IL-2, TGF-β and IL-10 were synthesized as a polycistronic gBlock (Genscript) and cloned into the existing GAPluc targeting plasmid as described above to generate the GAP-2B10 plasmid. The human IL-2 sequence was modified by substituting a single amino acid (N88D) as previously described (Peterson et al., 2018). GAP-2B10 hESCs were generated by co-nucleofection of the GAP-2B10 targeting plasmid and the GAPDH-targeting RNP as described above. GAP-2B10 SC-islet cells were differentiated as previously described (Millman et al., 2016; Pagliuca et al., 2014).

### In vitro glucose-stimulated insulin secretion

*In vitro* function was assessed by measuring glucose-stimulated insulin secretion (GSIS) as previously described (Millman et al., 2016; Pagliuca et al., 2014). SC-islet clusters were washed twice in Krebs buffer (KRB), and preincubated at 37 °C for 1 h in KRB containing 2 mM glucose (low glucose). Clusters were then challenged with three sequential treatments of alternating low-high-low KRB containing glucose (high; 20 mM), followed by depolarization with low KRB containing 30 mM KCl. Each treatment lasted 30 min, after which 100 μl of supernatant was collected and human insulin quantified using the Human Ultrasensitive Insulin ELISA. Human insulin measurements were normalized by viable cell counts that were acquired by dispersing clusters with TrypLE Express (Thermofisher, 12604013) and counted using a ViCell (Beckman Coulter).

### In vitro SC-islet cell cytokine secretion and PBMC cytotoxicity assays

GAP-2B10 SC-islet cell clusters were dispersed with TrypLE Express and seeded in 96-well matrigel-coated plates in S3 media at a linear concentration (1, 2, 4, 6, 8 x 10^4^, and 1 x 10^5^ cells/well) in duplicate. After 24 h, supernatants were collected, centrifuged at 3000g for 5 min and the cytokines IL-2, TGF-β and IL-10 quantified using Legend Max ELISAs (IL-2, Biolegend, 431807; TGF-β, Biolegend, 436707; IL-10, Biolegend, 430607) according to the manufacturer’s instructions. For cytotoxicity assays, WT and 2B10 SC-islet cells were co-cultured with human PBMCs as described above.

### Xenotransplantation of GAP-2B10 SC-islet cells

WT and GAP-2B10 SC-islet cell clusters (5 x 10^6^ cells) were transplanted under the kidney capsule of B6/albino mice (*n* = 3/group) and graft survival was monitored weekly for nine weeks by bioluminescence following i.p. injection of D-luciferin as described above. At five weeks post-transplantation SC-islet grafts were removed for immunohistochemical analysis of surviving INS^+^ cells and CD8^+^ T and FOXP3^+^ Treg cells.

### Immunohistochemistry of SC-islet grafts

The kidneys of B6/Albino mice transplanted with WT and 2B10 SC-islets were excised, fixed overnight in 4% paraformaldehyde (PFA) at room temperature and embedded in paraffin. Sections were pre-cleared with Histo-Clear, rehydrated using an ethanol gradient and antigen fixed by incubating in boiling antigen retrieval reagent (10 mM sodium citrate, pH 6.0) for 50 min. Slides were then blocked in 5% donkey serum for 1h and stained with Guinea pig anti-human Insulin (DAKO, A0564), Rat anti-mouse CD8α (Biolegend, 100702) and Mouse anti-mouse FOXP3 (Biolegend, 320002) overnight at 4 °C. The slides were then washed three times, incubated in secondary antibodies goat anti-mouse 594 (Life Technologies, A-11032), goat anti-rat 488 (Life Technologies, A-11006) and goat anti-guinea pig 647 (Life technologies, A-21450) for 2 h at room temperature, washed, mounted in Vectashield with DAPI (Vector Laboratories; H-1200), covered with coverslips and sealed with clear nail polish. Representative regions were imaged using Zeiss.Z2 with Apotome microscope. All primary and secondary antibodies were used at dilution of 1:200 and 1:500 respectively.

### Statistics

All data are presented as means ± SD and were analyzed by GraphPad Prism 9 (GraphPad Software). Statistically significant differences were determined either by one-way or two-way ANOVA, with Tukey’s and Sidak’s post-hoc test for multiple comparisons, and two-tailed t test for pairwise comparisons. p values are indicated in the figures as *p < 0.05, **p < 0.01, ***p < 0.005 and ****p < 0.001.

## Acknowledgements

D.A.M. is an investigator of the Howard Hughes Medical Institute. This work was supported by grants from the Harvard Stem Cell Institute (DP-0180-18-02), JDRF (5-COE-2020-967-M-N), and the JPB Foundation (Award #1094). We would like to thank Ramona Pop for discussions on manuscript content.

---

**Figure S1.**
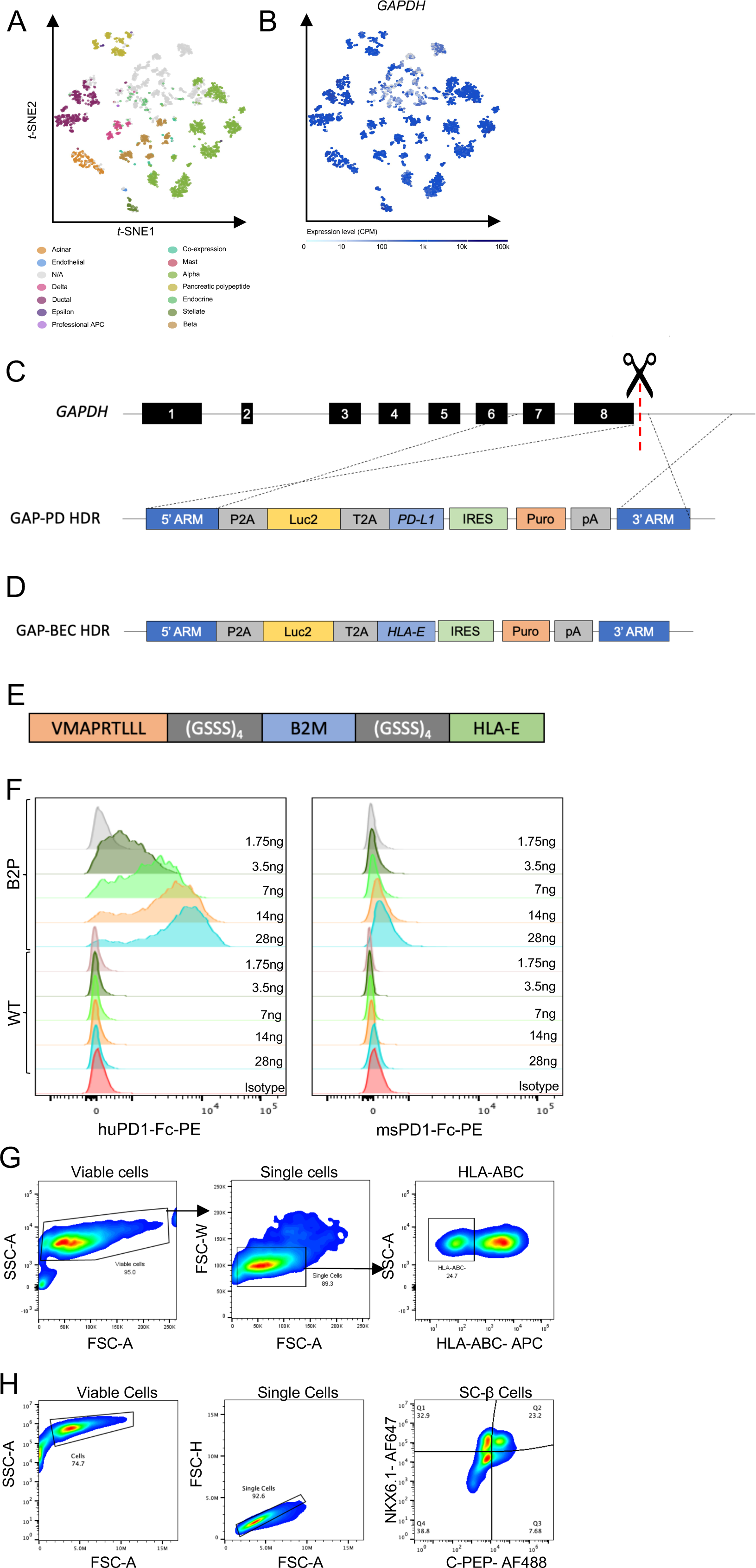
A. t-SNE projections of primary human islet cells. B. t-SNE projections of GAPDH expression across the assigned populations. Cells are colored according to their assigned cluster. (Adapted from Segerstolpe et al., 2016) C. Schematic of homology directed repair plasmids for integration of Luc2 and PD-L1 at the GAPDH locus. D. Schematic GAPDH-targeting Luc2 and peptide::B2M::HLA-E HDR plasmid. E. Schematic of the peptide::B2M::HLA-E long-chain fusion. F. PD1-Fc binding assay. PD-L1 and WT SC-islet cells were dissociated and stained with a 2-fold dilution series of PE-conjugated human and mouse PD1-Fc. Data is presented as MFI normalized to mode. G. FACS sorting strategy of HLA-ABC^-/-^ hESCs. H. FACS gating strategy for quantification of Nkx6.1^+^/C-pep^+^ SC-β cells following *in vitro* differentiation.

**Figure S2.**
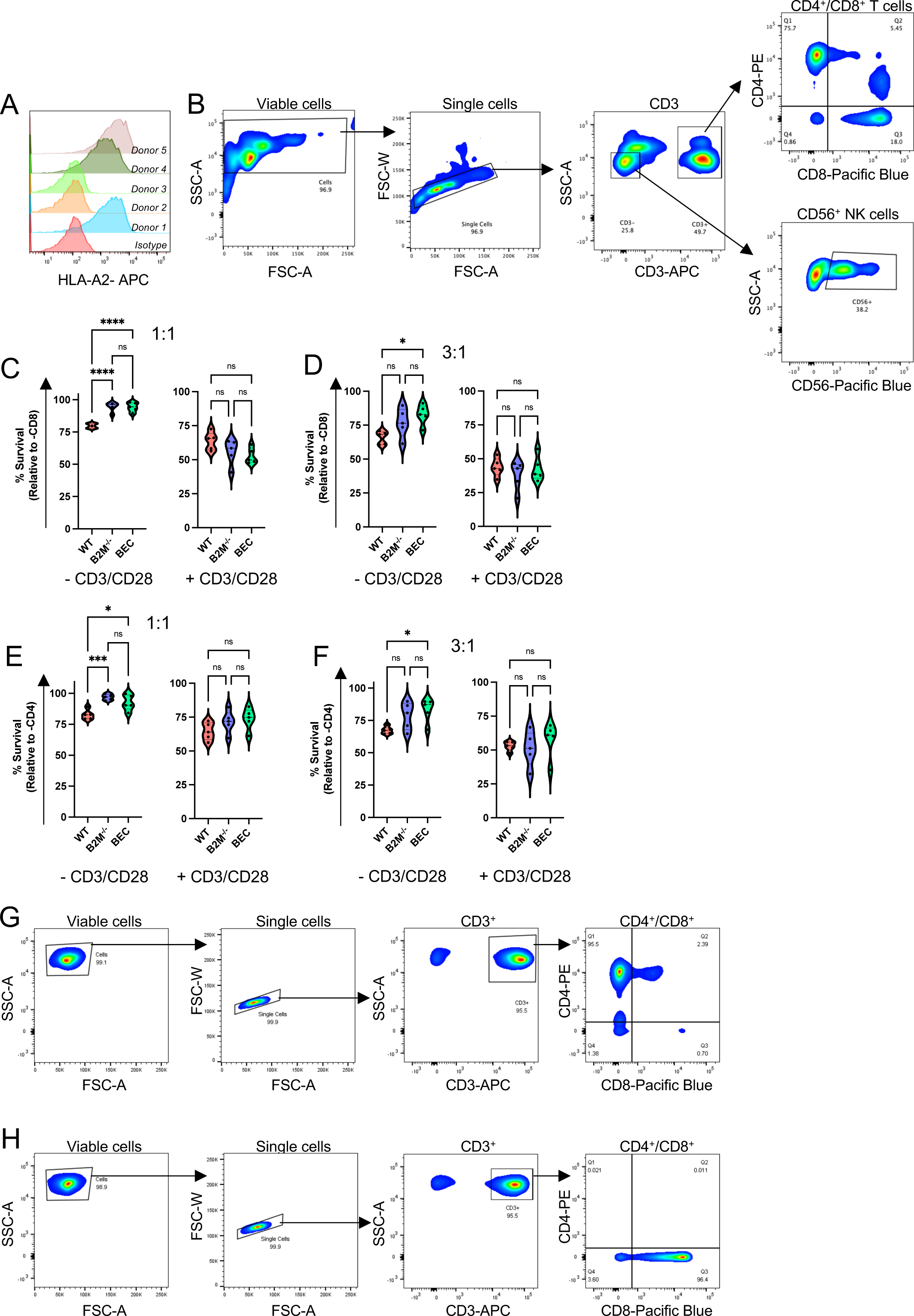
A. FACS profiling of HLA-A2 status of 5 PBMC donors. B. FACS gating strategy of CD4^+^ and CD8^+^ T cells and CD56^+^ NK cells in PBMCs enriched from human apheresis leukoreductions. Plots are representative of 5 donors. C. Quantification of SC-islet cell survival when co-cultured with purified human CD8^+^ T cells at a 1:1 ratio. Cell survival is presented as mean ± SD (*n* = 5). D. Quantification of SC-islet cell survival when co-cultured with purified human CD8^+^ T cells at a 3:1 ratio. Cell survival is presented as mean ± SD (*n* = 5). E. Quantification of SC-islet cell survival when co-cultured with purified human CD4^+^ T cells at a 1:1 ratio. Cell survival is presented as mean ± SD (*n* = 5). F. Quantification of SC-islet cell survival when co-cultured with purified human CD4^+^ T cells at a 3:1 ratio. Cell survival is presented as mean ± SD (*n* = 5). G. FACS gating strategy of CD4^+^ T cells enriched from human apheresis leukoreductions. Plots are representative of 5 donors. H. FACS gating strategy of CD8^+^ T cells enriched from human apheresis leukoreductions. Plots are representative of 5 donors.

**Figure S3.**
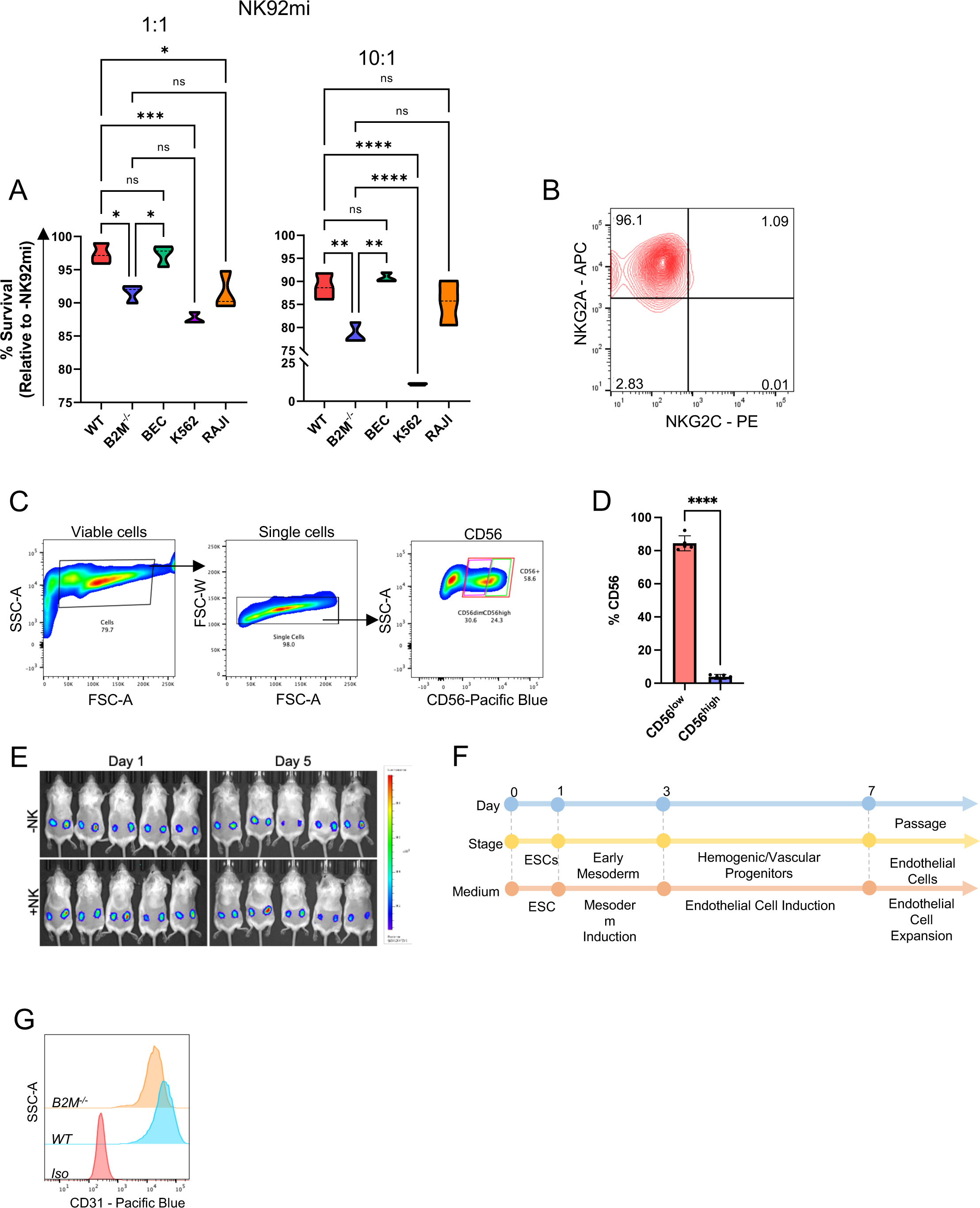
A. Quantification of SC-β cell survival when co-cultured with NK92mi cells. K562 and Raji cells were used as positive and negative controls, respectively. Cell survival is presented as mean ± SD (n = 5). B. NKG2A/NKG2C expression on NK92mi cells. C. FACS gating strategy of CD56^+^ NK cells enriched from human apheresis leukoreductions. Gates for specific population include all CD56^+^ NK cells (red), CD56^high^ (green) and CD56^dim^ (pink). Plots are representative of 5 donors. D. Quantitative analysis of CD56 expression on enriched primary human NK cells. Data is presented as % CD56 expression (*n* = 5). E. *In vivo* NK cell assay. Bioluminescence imaging was performed on day 1 and 5 post-transplantation. F. Schematic of SC-endothelial cell differentiation protocol. G. FACS analysis of the endothelial cell marker CD31 in SC-endothelial cells derived from WT and B2M^-/-^ hESCs.

## References

1. Abou-Daya, K.I., Tieu, R., Zhao, D., Rammal, R., Sacirbegovic, F., Williams, A.L., Shlomchik, W.D., Oberbarnscheidt, M.H., and Lakkis, F.G. (2021). Resident memory T cells form during persistent antigen exposure leading to allograft rejection. Science Immunology 6, eabc8122.

2. Alagpulinsa, D.A., Cao, J.J.L., Driscoll, R.K., Sîrbulescu, R.F., Penson, M.F.E., Sremac, M., Engquist, E.N., Brauns, T.A., Markmann, J.F., Melton, D.A., et al. (2019). Alginate-microencapsulation of human stem cell-derived β cells with CXCL12 prolongs their survival and function in immunocompetent mice without systemic immunosuppression. Am J Transplant 19, 1930–1940.

3. Atkinson, M.A., and Maclaren, N.K. (1994). The Pathogenesis of Insulin-Dependent Diabetes Mellitus. The New England Journal of Medicine 19, 1428–1436.

4. Benci, J.L., Xu, B., Qiu, Y., Wu, T.J., Dada, H., Twyman-Saint Victor, C., Cucolo, L., Lee, D.S.M., Pauken, K.E., Huang, A.C., et al. (2016). Tumor Interferon Signaling Regulates a Multigenic Resistance Program to Immune Checkpoint Blockade. Cell 167, 1540–1554.

5. Bochenek, M.A., Veiseh, O., Vegas, A.J., McGarrigle, J.J., Qi, M., Marchese, E., Omami, M., Doloff, J.C., Mendoza-Elias, J., Nourmohammadzadeh, M., et al. (2018). Alginate encapsulation as long-term immune protection of allogeneic pancreatic islet cells transplanted into the omental bursa of macaques. Nat Biomed Eng 2, 810–821.

6. Castro-Gutierrez, R., Alkanani, A., Mathews, C.E., Michels, A., and Russ, H.A. (2021). Protecting Stem Cell Derived Pancreatic Beta-Like Cells From Diabetogenic T Cell Recognition. Frontiers in Endocrinology 12.

7. Chen, L., and Flies, D.B. (2013). Molecular mechanisms of T cell co-stimulation and co-inhibition. Nat Rev Immunol 13, 227-242.

8. D’Amour, K.A., Bang, A.G., Eliazer, S., Kelly, O.G., Agulnick, A.D., Smart, N.G., Moorman, M.A., Kroon, E., Carpenter, M.K., and Baetge, E.E. (2006). Production of pancreatic hormone-expressing endocrine cells from human embryonic stem cells. Nat Biotechnol 24, 1392–1401.

9. Deuse, T., Hu, X., Agbor-Enoh, S., Jang, M.K., Alawi, M., Saygi, C., Gravina, A., Tediashvili, G., Nguyen, V.Q., Liu, Y., et al. (2021). The SIRPα-CD47 immune checkpoint in NK cells. The Journal of experimental medicine 218.

10. Deuse, T., Hu, X., Gravina, A., Wang, D., Tediashvili, G., De, C., Thayer, W.O., Wahl, A., Garcia, J.V., Reichenspurner, H., et al. (2019). Hypoimmunogenic derivatives of induced pluripotent stem cells evade immune rejection in fully immunocompetent allogeneic recipients. Nature Biotechnology 37, 252–258.

11. Finger, L.R., Pu, J., Wasserman, R., Vibhakar, R., Louie, E., Hardy, R.R., Burrows, P.D., and Billips, L.G. (1997). The human PD-1 gene: complete cDNA, genomic organization, and developmentally regulated expression in B cell progenitors. Gene 197, 177-187.

12. Freeman, G.J., Long, A.J., Iwai, Y., Bourque, K., Chernova, T., Nishimura, H., Fitz, L.J., Malenkovich, N., Okazaki, T., Byrne, M.C., et al. (2000). Engagement of the PD-1 immunoinhibitory receptor by a novel B7 family member leads to negative regulation of lymphocyte activation. The Journal of experimental medicine 192, 1027–1034.

13. Furman, B.L. (2021). Streptozotocin-Induced Diabetic Models in Mice and Rats. Current Protocols 1, e78.

14. Garcia-Diaz, A., Shin, D.S., Moreno, B.H., Saco, J., Escuin-Ordinas, H., Rodriguez, G.A., Zaretsky, J.M., Sun, L., Hugo, W., Wang, X., et al. (2017). Interferon Receptor Signaling Pathways Regulating PD-L1 and PD-L2 Expression. Cell reports 19, 1189-1201.

15. Gerace, D., Boulanger, K.R., Hyoje-Ryu Kenty, J., and Melton, D.A. (2021). Generation of a heterozygous GAPDH-Luciferase human ESC line (HVRDe008-A-1) for in vivo monitoring of stem cells and their differentiated progeny. Stem Cell Research 53, 102371.

16. Gornalusse, G.G., Hirata, R.K., Funk, S.E., Riolobos, L., Lopes, V.S., Manske, G., Prunkard, D., Colunga, A.G., Hanafi, L.-A., Clegg, D.O., et al. (2017). HLA-E-expressing pluripotent stem cells escape allogeneic responses and lysis by NK cells. Nature Biotechnology 35, 765.

17. Han, X., Wang, M., Duan, S., Franco, P.J., Kenty, J.H.-R., Hedrick, P., Xia, Y., Allen, A., Ferreira, L.M.R., Strominger, J.L., et al. (2019). Generation of hypoimmunogenic human pluripotent stem cells. Proceedings of the National Academy of Sciences 116, 10441.

18. Harding, J., Vintersten-Nagy, K., Shutova, M., Yang, H., Tang, J.K., Massumi, M., Izaidfar, M., Izadifar, Z., Zhang, P., Li, C., et al. (2019). Induction of long-term allogeneic cell acceptance and formation of immune privileged tissue in immunocompetent hosts. bioRxiv, 716571.

19. Hartemann, A., Bensimon, G., Payan, C.A., Jacqueminet, S., Bourron, O., Nicolas, N., Fonfrede, M., Rosenzwajg, M., Bernard, C., and Klatzmann, D. (2013). Low-dose interleukin 2 in patients with type 1 diabetes: a phase 1/2 randomised, double-blind, placebo-controlled trial. Lancet Diabetes Endocrinol 1, 295–305.

20. Herbst, F., Ball, C.R., Tuorto, F., Nowrouzi, A., Wang, W., Zavidij, O., Dieter, S.M., Fessler, S., van der Hoeven, F., Kloz, U., et al. (2012). Extensive Methylation of Promoter Sequences Silences Lentiviral Transgene Expression During Stem Cell Differentiation In Vivo. Molecular Therapy 20, 1014–1021.

21. Herndler-Brandstetter, D., Shan, L., Yao, Y., Stecher, C., Plajer, V., Lietzenmayer, M., Strowig, T., de Zoete, M.R., Palm, N.W., Chen, J., et al. (2017). Humanized mouse model supports development, function, and tissue residency of human natural killer cells. Proceedings of the National Academy of Sciences 114, E9626.

22. Herold, K.C., Bundy, B.N., Long, S.A., Bluestone, J.A., DiMeglio, L.A., Dufort, M.J., Gitelman, S.E., Gottlieb, P.A., Krischer, J.P., Linsley, P.S., et al. (2019). An Anti-CD3 Antibody, Teplizumab, in Relatives at Risk for Type 1 Diabetes. New England Journal of Medicine 381, 603-613.

23. Heusschen, R., Griffioen, A.W., and Thijssen, V.L. (2013). Galectin-9 in tumor biology: A jack of multiple trades. Biochimica et Biophysica Acta (BBA) - Reviews on Cancer 1836, 177–185.

24. Horwitz, D.A., Zheng, S.G., Wang, J., and Gray, J.D. (2008). Critical role of IL-2 and TGF-beta in generation, function and stabilization of Foxp3+CD4+ Treg. Eur J Immunol 38, 912–915.

25. Imaizumi, T., Kumagai, M., Sasaki, N., Kurotaki, H., Mori, F., Seki, M., Nishi, N., Fujimoto, K., Tanji, K., Shibata, T., et al. (2002). Interferon-gamma stimulates the expression of galectin-9 in cultured human endothelial cells. J Leukoc Biol 72, 486–491.

26. Kaiser, B.K., Barahmand-pour, F., Paulsene, W., Medley, S., Geraghty, D.E., and Strong, R.K. (2005). Interactions between NKG2x Immunoreceptors and HLA-E Ligands Display Overlapping Affinities and Thermodynamics. The Journal of Immunology 174, 2878.

27. Khoryati, L., Pham, M.N., Sherve, M., Kumari, S., Cook, K., Pearson, J., Bogdani, M., Campbell, D.J., and Gavin, M.A. (2020). An IL-2 mutein engineered to promote expansion of regulatory T cells arrests ongoing autoimmunity in mice. Science immunology 5, eaba5264.

28. Kleiveland, C.R. (2015). Peripheral Blood Mononuclear Cells. In The Impact of Food Bioactives on Health: in vitro and ex vivo models, K. Verhoeckx, P. Cotter, I. López-Expósito, C. Kleiveland, T. Lea, A. Mackie, T. Requena, D. Swiatecka, and H. Wichers, eds. (Cham: Springer International Publishing), pp. 161–167.

29. Leite, N.C., Pelayo, G.C., and Melton, D.A. (2022). Genetic manipulation of stress pathways can protect stem-cell-derived islets from apoptosis in vitro. Stem cell reports.

30. Lim, D., Sreekanth, V., Cox, K.J., Law, B.K., Wagner, B.K., Karp, J.M., and Choudhary, A. (2020). Engineering designer beta cells with a CRISPR-Cas9 conjugation platform. Nature communications 11, 4043.

31. Mandal, P.K., Ferreira, L.M., Collins, R., Meissner, T.B., Boutwell, C.L., Friesen, M., Vrbanac, V., Garrison, B.S., Stortchevoi, A., Bryder, D., et al. (2014). Efficient ablation of genes in human hematopoietic stem and effector cells using CRISPR/Cas9. Cell stem cell 15, 643–652.

32. Mattapally, S., Pawlik, K.M., Fast, V.G., Zumaquero, E., Lund, F.E., Randall, T.D., Townes, T.M., and Zhang, J. (2018). Human Leukocyte Antigen Class I and II Knockout Human Induced Pluripotent Stem Cell-Derived Cells: Universal Donor for Cell Therapy. J Am Heart Assoc 7, e010239.

33. Millman, J.R., Xie, C., Van Dervort, A., Gurtler, M., Pagliuca, F.W., and Melton, D.A. (2016). Generation of stem cell-derived beta-cells from patients with type 1 diabetes. Nature communications 7, 11463.

34. Monti, P., Scirpoli, M., Maffi, P., Ghidoli, N., De Taddeo, F., Bertuzzi, F., Piemonti, L., Falcone, M., Secchi, A., and Bonifacio, E. (2008). Islet transplantation in patients with autoimmune diabetes induces homeostatic cytokines that expand autoreactive memory T cells. The Journal of clinical investigation 118, 1806–1814.

35. Nair, G.G., Liu, J.S., Russ, H.A., Tran, S., Saxton, M.S., Chen, R., Juang, C., Li, M.L., Nguyen, V.Q., Giacometti, S., et al. (2019). Recapitulating endocrine cell clustering in culture promotes maturation of human stem-cell-derived beta cells. Nature cell biology 21, 263–274.

36. Oberbarnscheidt, M.H., Zeng, Q., Li, Q., Dai, H., Williams, A.L., Shlomchik, W.D., Rothstein, D.M., and Lakkis, F.G. (2014). Non-self recognition by monocytes initiates allograft rejection. The Journal of clinical investigation 124, 3579–3589.

37. Pagliuca, F.W., Millman, J.R., Gurtler, M., Segel, M., Van Dervort, A., Ryu, J.H., Peterson, Q.P., Greiner, D., and Melton, D.A. (2014). Generation of functional human pancreatic beta cells in vitro. Cell 159, 428–439.

38. Parent, A.V., Faleo, G., Chavez, J., Saxton, M., Berrios, D.I., Kerper, N.R., Tang, Q., and Hebrok, M. (2021). Selective deletion of human leukocyte antigens protects stem cell-derived islets from immune rejection. Cell Reports 36, 109538.

39. Peterson, L.B., Bell, C.J.M., Howlett, S.K., Pekalski, M.L., Brady, K., Hinton, H., Sauter, D., Todd, J.A., Umana, P., Ast, O., et al. (2018). A long-lived IL-2 mutein that selectively activates and expands regulatory T cells as a therapy for autoimmune disease. Journal of Autoimmunity 95, 1–14.

40. Raudvere, U., Kolberg, L., Kuzmin, I., Arak, T., Adler, P., Peterson, H., and Vilo, J. (2019). g:Profiler: a web server for functional enrichment analysis and conversions of gene lists (2019 update). Nucleic acids research 47, W191–W198.

41. Rezania, A., Bruin, J.E., Arora, P., Rubin, A., Batushansky, I., Asadi, A., O’Dwyer, S., Quiskamp, N., Mojibian, M., Albrecht, T., et al. (2014). Reversal of diabetes with insulin-producing cells derived in vitro from human pluripotent stem cells. Nat Biotechnol 32, 1121–1133.

42. Rigau, M., Ostrouska, S., Fulford Thomas, S., Johnson Darryl, N., Woods, K., Ruan, Z., McWilliam Hamish, E.G., Hudson, C., Tutuka, C., Wheatley Adam, K., et al. (2020). Butyrophilin 2A1 is essential for phosphoantigen reactivity by γδ T cells. Science 367, eaay5516.

43. Riolobos, L., Hirata, R.K., Turtle, C.J., Wang, P.R., Gornalusse, G.G., Zavajlevski, M., Riddell, S.R., and Russell, D.W. (2013). HLA engineering of human pluripotent stem cells. Molecular therapy : the journal of the American Society of Gene Therapy 21, 1232–1241.

44. Russ, H.A., Parent, A.V., Ringler, J.J., Hennings, T.G., Nair, G.G., Shveygert, M., Guo, T., Puri, S., Haataja, L., Cirulli, V., et al. (2015). Controlled induction of human pancreatic progenitors produces functional beta-like cells in vitro. The EMBO journal 34, 1759–1772.

45. Shapiro, A.M., Ricordi, C., Hering, B.J., Auchincloss, H., Lindblad, R., Robertson, R.P., Secchi, A., Brendel, M.D., Berney, T., Brennan, D.C., et al. (2006). International trial of the Edmonton protocol for islet transplantation. N Engl J Med 355, 1318–1330.

46. Sintov, E., Gerace, D., and Melton, D.A. (2021). A human ESC line for efficient CRISPR editing of pluripotent stem cells. Stem Cell Research 57, 102591.

47. Stegall, M.D., Lafferty, K.J., Kam, I., and Gill, R.G. (1996). Evidence of recurrent autoimmunity in human allogeneic islet transplantation. Transplantation 61, 1272–1274.

48. Suzuki, D., Flahou, C., Yoshikawa, N., Stirblyte, I., Hayashi, Y., Sawaguchi, A., Akasaka, M., Nakamura, S., Higashi, N., Xu, H., et al. (2020). iPSC-Derived Platelets Depleted of HLA Class I Are Inert to Anti-HLA Class I and Natural Killer Cell Immunity. Stem cell reports 14, 49–59.

49. Veres, A., Faust, A.L., Bushnell, H.L., Engquist, E.N., Kenty, J.H., Harb, G., Poh, Y.C., Sintov, E., Gurtler, M., Pagliuca, F.W., et al. (2019). Charting cellular identity during human in vitro beta-cell differentiation. Nature 569, 368–373.

50. Viricel, C., Ahmed, M., and Barakat, K. (2015). Human PD-1 binds differently to its human ligands: a comprehensive modeling study. J Mol Graph Model 57, 131–142.

51. Wagner, A.H., Gebauer, M., Pollok-Kopp, B., and Hecker, M. (2002). Cytokine-inducible CD40 expression in human endothelial cells is mediated by interferon regulatory factor-1. Blood 99, 520–525.

52. Wang, D., Quan, Y., Yan, Q., Morales, J.E., and Wetsel, R.A. (2015). Targeted Disruption of the β2-Microglobulin Gene Minimizes the Immunogenicity of Human Embryonic Stem Cells. Stem cells translational medicine 4, 1234–1245.

53. Wen, J., Wu, J., Cao, T., Zhi, S., Chen, Y., Aagaard, L., Zhen, P., Huang, Y., Zhong, J., and Huang, J. (2021). Methylation silencing and reactivation of exogenous genes in lentivirus-mediated transgenic mice. Transgenic Res 30, 63–76.

54. Wyburn, K.R., Jose, M.D., Wu, H., Atkins, R.C., and Chadban, S.J. (2005). The Role of Macrophages in Allograft Rejection. Transplantation 80.

55. Xu, H., Wang, B., Ono, M., Kagita, A., Fujii, K., Sasakawa, N., Ueda, T., Gee, P., Nishikawa, M., Nomura, M., et al. (2019). Targeted Disruption of HLA Genes via CRISPR-Cas9 Generates iPSCs with Enhanced Immune Compatibility. Cell stem cell 24, 566–578.e567.

56. Yang, R., Sun, L., Li, C.-F., Wang, Y.-H., Yao, J., Li, H., Yan, M., Chang, W.-C., Hsu, J.-M., Cha, J.-H., et al. (2021). Galectin-9 interacts with PD-1 and TIM-3 to regulate T cell death and is a target for cancer immunotherapy. Nature communications 12, 832.

57. Yoshihara, E., O’Connor, C., Gasser, E., Wei, Z., Oh, T.G., Tseng, T.W., Wang, D., Cayabyab, F., Dai, Y., Yu, R.T., et al. (2020). Immune-evasive human islet-like organoids ameliorate diabetes. Nature 586, 606–611.

58. Zhuang, Q., Liu, Q., Divito, S.J., Zeng, Q., Yatim, K.M., Hughes, A.D., Rojas-Canales, D.M., Nakao, A., Shufesky, W.J., Williams, A.L., et al. (2016). Graft-infiltrating host dendritic cells play a key role in organ transplant rejection. Nature communications 7, 12623-12623.

